# Immunity induced by vaccination with recombinant influenza B virus neuraminidase protein breaks viral transmission chains in guinea pigs in an exposure intensity-dependent manner

**DOI:** 10.1101/2022.10.19.512980

**Authors:** Meagan McMahon, Jessica Tan, George O’Dell, Ericka Kirkpatrick Roubidoux, Shirin Strohmeier, Florian Krammer

## Abstract

Mucosal vaccines and vaccines that block pathogen transmission are under-appreciated in vaccine development. However, the severe acute respiratory syndrome coronavirus 2 (SARS-CoV-2) pandemic has shown that blocking viral transmission is an important attribute of efficient vaccines. Here, we investigated if recombinant influenza virus neuraminidase (NA) vaccines delivered at a mucosal site could protect from onward transmission of influenza B viruses in the guinea pig model. We tested four different scenarios in which sequential transmission was investigated in chains of four guinea pigs. The variables tested included a low and a high viral inoculum (10^4^ vs 10^5^ plaque forming units) in the initial donor guinea pig and variation of exposure/cohousing time (1 day vs 6 days). In three out of four scenarios – low inoculum-long exposure, low inoculum-short exposure and high inoculum-short exposure – transmission chains were efficiently blocked. Based on this data we believe an intranasal recombinant NA vaccine could be used to efficiently curtail influenza virus spread in the human population during influenza epidemics.

**Importance:** Vaccines that can slow respiratory virus transmission in the population are urgently needed for severe acute respiratory syndrome coronavirus 2 (SARS-CoV-2) and influenza virus. Here we describe how a recombinant neuraminidase-based influenza virus vaccines reduces transmission in vaccinated guinea pigs in an exposure-intensity based manner.

## Introduction

The current severe acute respiratory syndrome coronavirus 2 (SARS-CoV-2) pandemic has highlighted how important it is that vaccines not only protect from disease but also limit onward transmission of pathogens. Similar to injected SARS-CoV-2 vaccines, influenza virus vaccines – even if well-matched – often allow onward transmission of virus in vaccinated populations (1). The intramuscular administration of current SARS-CoV-2 vaccines as well as inactivated influenza virus vaccines contributes to this problem since this route of administration does not lead to robust mucosal antibody titers which could block infection or limit transmission (2-4).

Influenza virus vaccines typically induce an immune response focused on the viral hemagglutinin (HA), the receptor binding protein of influenza viruses which binds to terminal sialic acid on N-linked glycans on host cells. Immunity to HA can neutralize virus efficiently and block infection. However, the location of the vaccine-induced antibodies in combination with the constant changes of the HA through antigenic drift often lead to suboptimal immunity after vaccination. Besides HA, influenza viruses express a second surface glycoprotein, the neuraminidase (NA), which is a receptor destroying enzyme that cleaves terminal sialic acids from N-linked glycans (5-7). This activity is important for migration of incoming virus through mucosal fluids (8, 9). Mucosal fluids have high concentrations of glycosylated natural defense proteins, which can act as a virus trap to prevent the release of newly formed viral particles from infected cells. The presence of NA enzymatic activity releases cell surface bound virus and counters virus aggregation (10).

While both HA and NA proteins undergo antigenic drift, their drift is usually discordant and NA potentially evolves more slowly (11, 12). This, combined with its important function in the viral life cycle, makes it an attractive vaccine target. We and others have shown that vaccination with recombinant, stabilized NA protein can induce protective immunity in different animal models, especially when the antigen is given mucosally (13-21). Of note, this protection is typically against morbidity and mortality, and while viral replication in animal models is reduced, NA-based immunity is often infection permissive. Here, we use the well established guinea pig influenza virus transmission model (22) to determine if vaccination with recombinant influenza B virus neuraminidase can break viral transmission chains and which factors may influence efficiency of transmission in the background of mucosal NA immunity.

## Methods

### Viruses and cells

Sf9 cells (CRL-1711, ATCC) for baculovirus rescue were grown in *Trichoplusia ni* medium-formulation Hink insect cell medium (TNM-FH, Gemini Bioproducts) supplemented with 10% fetal bovine serum (FBS; Sigma) and penicillin (100 U/ml)-streptomycin (100 μg/ml) solution (Gibco). BTI-*TN*-5B1-4 (High Five, ATCC) cells for protein expression were grown in serum-free Express Five SFM media (Gibco) supplemented with penicillin (100 U/ml)-streptomycin (100 μg/ml) solution. Madin Darby canine kidney (MDCK, ATCC) cells were grown in Dulbecco’s modified Eagle’s medium (DMEM, Gibco) supplemented with 10% FBS and penicillin (100 U/ml)-streptomycin (100 μg/ml) solution. B/Malaysia/2506/04 virus was grown in 10-day-old embryonated chicken eggs (Charles River) for 72 hours at 33°C. Eggs were then cooled overnight at 4°C before harvesting the allantoic fluid. Harvested allantoic fluid was centrifuged at 4,000 g for 10 minutes at 4°C to pellet debris. Viruses were then aliquoted and stored at -80°C prior to determining stock titers via plaque assay.

### Protein production

Recombinant NAs from A/Michigan/45/15 (H1N1) or B/Malaysia/2506/04 virus were expressed in High Five insect cells as a fusion protein with an N-terminal vasodilator-stimulated phosphoprotein (VASP) tetramerization domain (23) and the globular head domain of the NA. Proteins were purified from the cell culture supernatant via Ni^2+^-nitrilotriacetic acid (Ni-NTA) chromatography (24, 25).

### Guinea pig vaccination

All animal experiments were conducted in concordance with protocols approved by the Icahn School of Medicine at Mount Sinai Institutional Animal Care and Use Committee. Five-to six-week-old female guinea pigs were purchased from Charles River Laboratory and randomly assigned to different vaccination groups. Guinea pigs were primed intranasally (I.N.) with 10 μg of A/Michigan/45/15 (N1) or B/Malaysia/2506/04 NA adjuvanted with 10 μg of poly(I⋅C) (Invivogen). Four weeks after the prime, a boost via the I.N. route with 10 μg of poly(I⋅C)-adjuvanted recombinant protein was administered. At 4 weeks post boost, vaccinated guinea pigs were used in transmission studies.

### Transmission experiments

Co-caged guinea pig transmission experiments were performed as previously described (26). For transmission studies where guinea pigs were co-caged with initial donors for 6 days **(Fig 1A)**, naïve donor guinea pigs were anaesthetized with ketamine (30 mg/kg) and xylazine (5 mg/kg) before being challenged I.N. with 10^4^ or 10^5^ plaque forming units (PFU) of B/Malaysia/2506/04 in 300 μL of phosphate-buffered saline (PBS). The following day, donor and vaccinated recipient (recipient 1) transmission pairs were co-caged (contact transmission). On day 6 post initial donor challenge, the recipient guinea pig (recipient 1) was removed and rehoused with another vaccinated recipient guinea pig (recipient 2). Recipient 2 was re-homed again on day 12 post initial donor challenge with vaccinated recipient 3. On days 2, 4, 6, 8, and 10 post contact, nasal washes were collected from anaesthetized donor and recipient guinea pigs. Recipient 2 guinea pigs received additional nasal washes on day 12 and 14 post contact.

**Figure 1.**
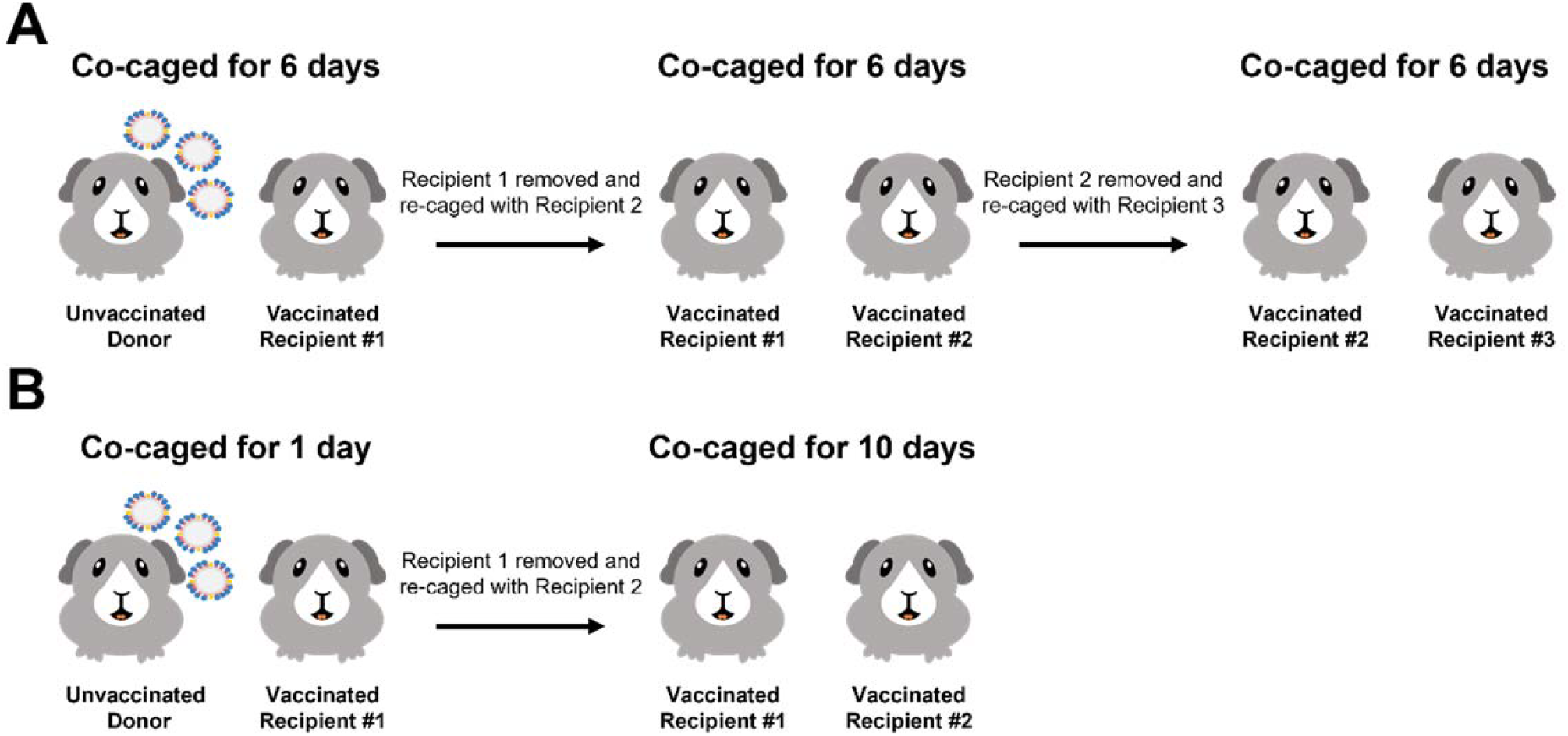
Schematic depicting the transmission settings used in these studies. The experimental setup for the 6-day contact transmission setting **(A)** and 1-day contact transmission setting **(B)**.

For transmission studies where guinea pigs were co-caged with initial donors for 1 day **(Fig 1B)**, naïve donor guinea pigs were anaesthetized and challenged as described above. The following day, donor and vaccinated recipient transmission pairs were co-caged (contact transmission). On the subsequent day, vaccinated recipient guinea pigs were removed and re-housed with another vaccinated recipient guinea pig. On days 2, 4, 6, 8, and 10 post contact, nasal washes were collected from anaesthetized donor and recipient guinea pigs.

### Plaque assays

Virus titers were determined by plaque assay on MDCK cell monolayers. Virus stocks and nasal washes were diluted 10-fold in 1× minimum essential medium (MEM) (10% 10× minimal essential medium [Gibco], 2 mM L-glutamine, 0.1% of sodium bicarbonate [wt/vol; Gibco], 10 mM 4-(2-hydroxyethyl)-1-piperazineethanesulfonic acid (HEPES) (Gibco), 100 U/ml penicillin–100 μ/ml streptomycin, and 0.2% bovine serum albumin (BSA) and 0.1% (wt/vol) diethylaminoethyl (DEAE)-dextran was added to the cells and incubated on MDCK cells for 1 hour before the an agarose overlay containing a final concentration of 0.64% agarose (Oxoid), 1x MEM and 1U/mL tolylsulfonyl phenylalanyl chloromethyl ketone (TPCK)-treated trypsin was added to the cells. The cells were then incubated for 72 hours at 33°C, and visible plaques were counted after fixation with 3.7% formaldehyde and visualization with a crystal violet counterstain (Sigma-Aldrich). All virus titers are presented as the log_10_ PFU/mL. The limit of detection for these assays was 50 PFU/mL.

## Results

### Intranasal vaccination with B/Malaysia/2506/2004 NA limits transmission between co-caged guinea pigs, although this is inoculation titer-dependent

Our previous work found that transmission from naïve B/Malaysia/2506/2004 infected donors to B/Malaysia/2506/2004 NA vaccinated recipients in a contact transmission setting results in transmission to three of three vaccinated recipients – meaning in that setting transmission was not prevented (26). However, we noted in this work that these vaccinated guinea pigs had very low nasal wash titers and a short duration of shedding. Here, we wanted to determine if these infected, but vaccinated guinea pigs, could allow subsequent infection.

In these studies we initially infected naïve donor guinea pigs with 10^4^ PFU of B/Malaysia/2506/2004 virus. The following day, donor guinea pigs were co-caged with A/Michigan/45/2015 N1 (negative control group, **Fig 2A**) or B/Malaysia/2506/2004 NA (**Fig 2B**) vaccinated guinea pigs (recipient 1). On day 6 following the initial donor infection, recipient 1 guinea pigs were co-caged with vaccinated guinea pigs (recipient 2). On day 12 following the initial donor infection, recipient 2 guinea pigs were co-caged with vaccinated guinea pigs (recipient 3). We assessed virus titers in the nasal washes at days 2, 4, 6, 8 and 10 post initial contact. Virus titers in the nasal wash indicate that virus was transmitted to each recipient in the irrelevant NA vaccinated guinea pigs but virus transmission did not progress past recipient 1 in the B/Malaysia/2506/2004 NA vaccinated guinea pigs.

**Figure 2.**
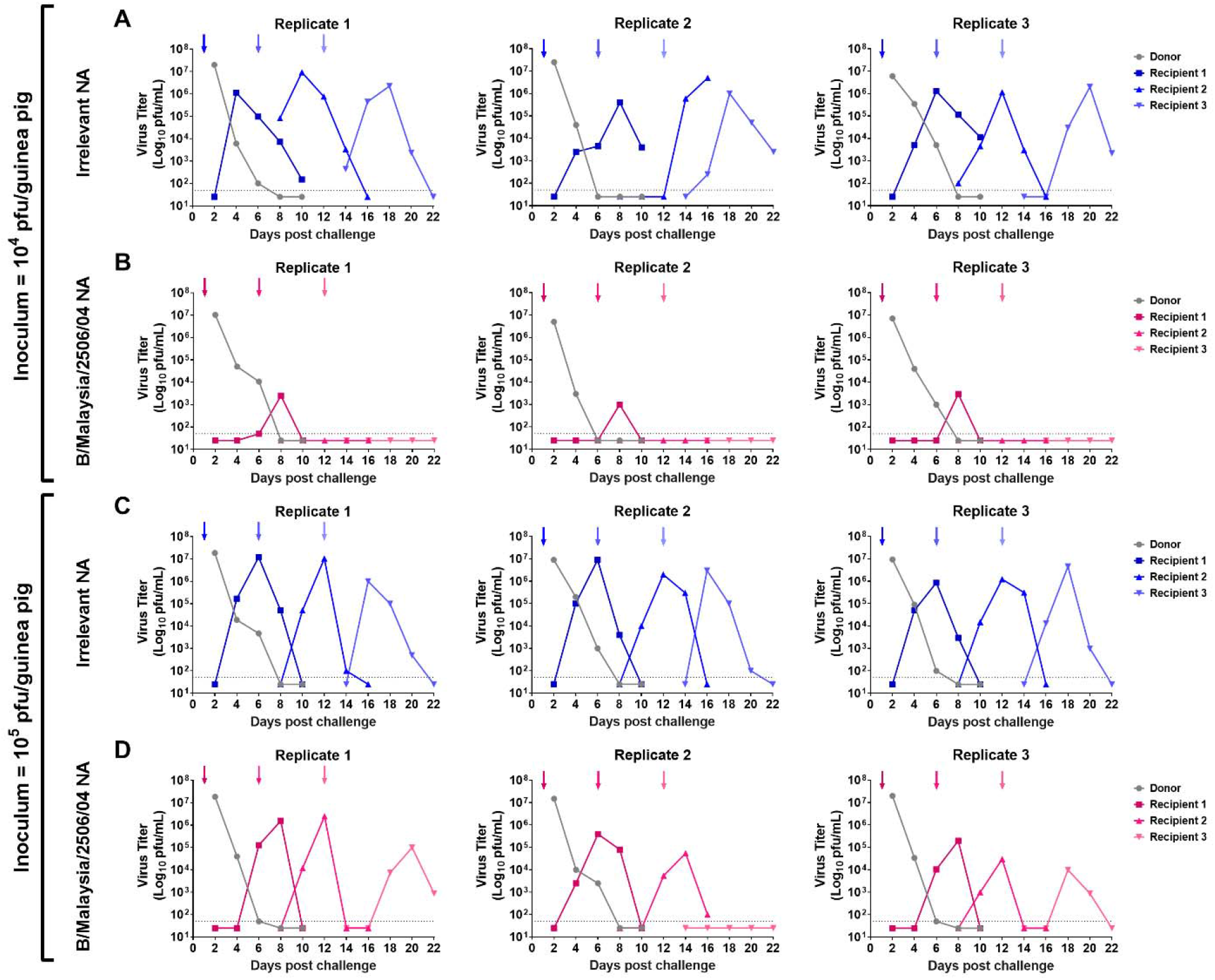
Assessment of B/Malaysia/2506/2004 transmission between vaccinated guinea pigs in a 6-day contact transmission setting. Naïve donor guinea pigs were anaesthetized and challenged with 10^4^ (A and B) or 10^5^ (C and D) PFU of B/Malaysia/2506/2004. The following day, donor and vaccinated recipient (recipient 1) transmission pairs were co-caged (contact transmission). On day 6 post initial donor challenge, the recipient guinea pig (recipient 1) was removed and rehoused with another vaccinated recipient guinea pig (recipient 2). Recipient 2 was re-homed again on day 12 post initial donor challenge with vaccinated recipient 3. On days 2, 4, 6, 8, and 10 post contact, nasal washes were collected from anaesthetized donor (gray) and recipient (non-gray) guinea pigs. Arrows depict the addition of a recipient and the removal of a donor/recipient. The experiment was repeated 3 times with each replicate containing an unvaccinated donor and a recipient 1, recipient 2 and recipient 3.

We next wanted to determine if increasing the inoculum titer would result in more efficient subsequent infection. Here we infected naïve donor guinea pigs with 10^5^ PFU of B/Malaysia/2506/2004 virus and performed recipient co-caging and nasal washes as described above. We found that, like above, virus transmitted to all of the irrelevant NA vaccinated guinea pigs (**Fig 2C**). Interestingly, we observed that virus transmitted from recipient 1 to recipient 2 in all of the B/Malaysia/2506/2004 NA replicates and virus transmitted from recipient 2 to recipient 3 in 2 out 3 B/Malaysia/2506/2004 NA.

These studies suggest that vaccination with B/Malaysia/2506/2004 NA is infection permissive, but subsequent transmission from NA vaccinated guinea pigs to other NA vaccinated guinea pigs can be blocked in a titer-dependent manner.

### Inhibition of transmission in recombinant NA vaccinated guinea pigs is dependent on the length of exposure to infected donor animals

After determining that B/Malaysia/2506/2004 NA vaccinated guinea pigs are susceptible to infection when exposed to infected guinea pigs for 6 days, we wanted to learn if B/Malaysia/2506/2004 NA vaccinated guinea pigs would be susceptible if exposure time is limited in duration (27). In these next experiments we infected naïve donor guinea pigs with 10^4^ (**Fig 3A and 3B**) or 10^5^ (**Fig 3C and 3D**) PFU of B/Malaysia/2506/2004 virus. The following day, donor guinea pigs were co-caged with A/Michigan/45/2015 N1 (negative control group, **Fig 3A or 3C**) or B/Malaysia/2506/2004 NA (**Fig 3B or 3D**) vaccinated guinea pigs (recipient 1). On day 2 following the initial donor infection, recipient 1 guinea pigs were co-caged with vaccinated guinea pigs (recipient 2) for the remainder of the experiment. We assessed virus titers in the nasal washes at days 2, 4, 6, 8 and 10 post donor infection in donors and recipient 1 and at days 4, 6, 8, 10, 12, 14 and 16 post donor infection for recipient 2 guinea pigs.

**Figure 3.**
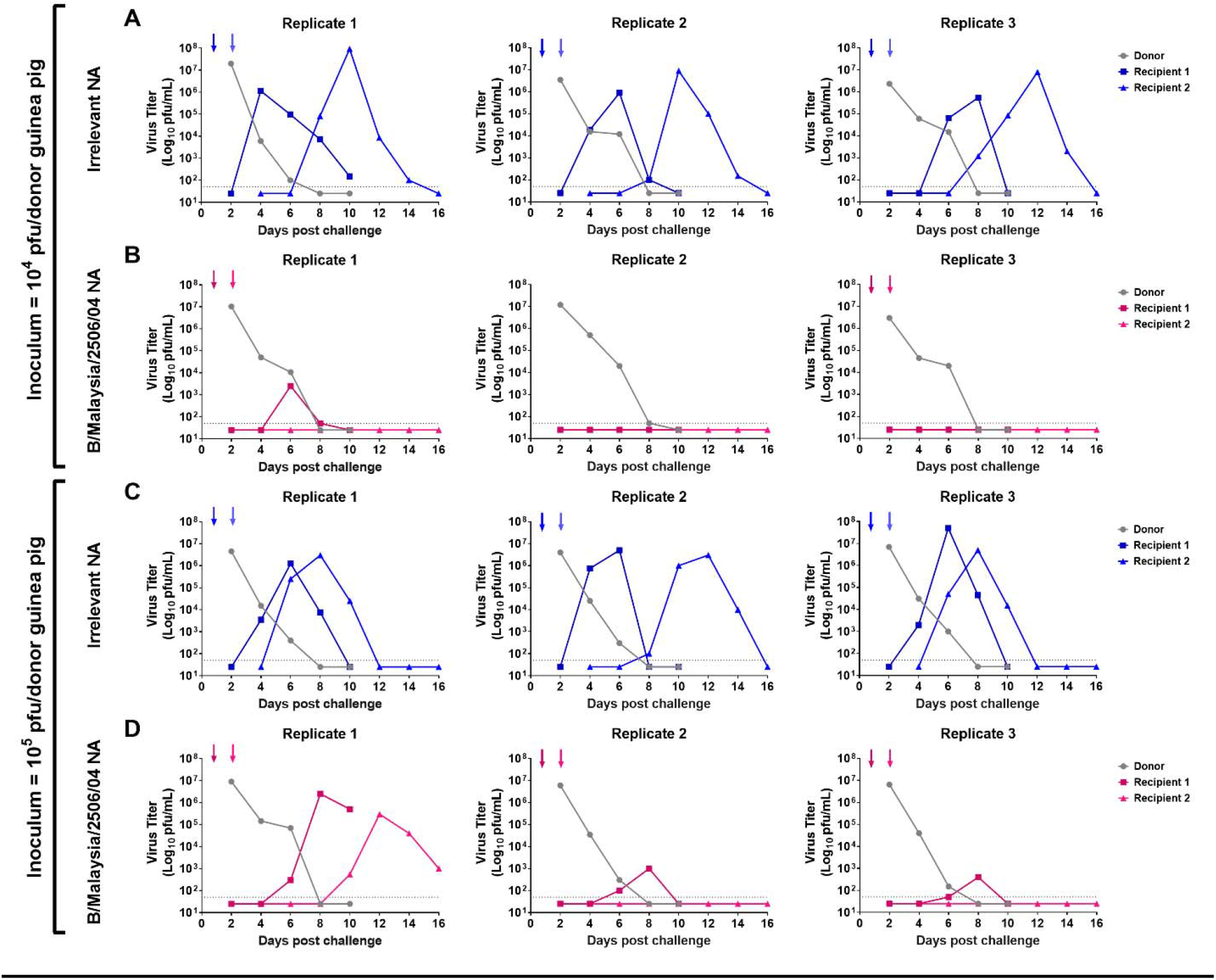
Assessment of B/Malaysia/2506/2004 transmission between vaccinated guinea pigs in a 1-day contact transmission setting. Naïve donor guinea pigs were anaesthetized and challenged with 10^4^ (A and B) or 10^5^ (C and D) PFU of B/Malaysia/2506/2004. The following day, donor and vaccinated recipient transmission pairs were co-caged. On the subsequent day, vaccinated recipient guinea pigs were removed and re-housed with another vaccinated recipient guinea pig. On days 2, 4, 6, 8, and 10 post contact, nasal washes were collected from anaesthetized donor (gray) and recipient (non-gray) guinea pigs. Recipient 2 guinea pigs received additional nasal washes on day 12 and 14 post contact. Arrows depict the addition of a recipient and the removal of a donor/recipient. The experiment was repeated 3 times with each replicate containing an unvaccinated donor and a recipient 1, recipient 2 and recipient 3.

In studies where naïve donor guinea pigs were infected with 10^4^ PFU of B/Malaysia/2506/2004, virus titer data indicate that virus was transmitted from the naïve donor to recipient 1 then on to recipient 2 in the irrelevant NA vaccinated guinea pigs in all 3 replicates (**Fig 3A**). In the B/Malaysia/2506/2004 NA vaccinated guinea pigs, virus was transmitted from the naïve donor to recipient 1 in only 1 out of 3 replicates (**Fig 3B**) and recipient 2 guinea pigs remained uninfected in all replicates.

In studies where naïve donor guinea pigs were infected with 10^5^ PFU of B/Malaysia/2506/2004 virus, virus titer data indicate that virus was transmitted from the naïve donor to recipient 1 then on to recipient 2 in the irrelevant NA vaccinated guinea pigs in all 3 replicates (**Fig 3C**). In the B/Malaysia/2506/2004 NA vaccinated guinea pigs, virus transmitted from the naïve donor to recipient 1 in all 3 replicates (**Fig 3B**), and from recipient 1 to recipient 2 in 1 out of 3 replicates.

These studies suggest that vaccination with B/Malaysia/2506/2004 NA, alongside relatively limited exposure to infected donors, resulted in reduced transmission to B/Malaysia/2506/2004 NA vaccinated guinea pigs.

## Discussion

Optimal vaccines serve two important purposes. They should protect the vaccinated individual from disease and they should protect others – including immunocompromised or naïve individuals – from onward transmission. While the second purpose was well recognized in the vaccinology and public health community, it has become part of public discourse during the SARS-CoV-2 pandemic. Several licensed vaccines fulfill both purposes. However, especially for respiratory viruses, blocking transmission through vaccination is often challenging as demonstrated with SARS-CoV-2 but also influenza virus. Part of the problem is that many vaccines are administered intramuscularly which makes them very inefficient in inducing mucosal immune responses (2-4). However, mucosal immune responses can block infection completely (sterilizing immunity) depending on the vaccine target, and they can blunt transmission by reducing titers and/or potentially by producing pathogen that is already coated in antibody when it leaves the upper respiratory tract and therefore perhaps reduce the infectiousness of an infected subject.

In the past we have shown that intranasal vaccination of guinea pigs, which are an excellent model for influenza virus transmission (while they do not show symptoms of disease), with recombinant NA can block viral transmission (18). However, this depended on the setting, and efficacy was higher in an ‘aerosol’ transmission setting in which animals were separated by perforated barriers as compared to a cohoused setting which allowed for direct contact. Interestingly, vaccinated guinea pigs, while supporting virus replication when directly infected, did not pass virus on to naïve animals (18). Vice versa, vaccinated guinea pigs exposed to naïve infected guinea pigs did get infected but experienced lower virus replication. Here, we wanted to investigate if NA vaccination could block transmission chains in a setting that previously led to greater transmission: directly cohousing vaccinated recipient animals with naïve infected donor animals. In this setting we wanted to explore two variables: Does virus dose of inoculation matter when initially infecting the donor guinea pig? And does the time donors and recipients are co-housed have an impact on transmission? We found that mucosal vaccination with recombinant NA can efficiently break transmission chains but this depends on ‘intensity’ of exposure. When donor animals were inoculated with a lower dose of virus and cohoused for a long period of time (6 days) with recipients, transmission to recipients occurred but only low viral titers were measured and virus was not further transmitted. If the initial viral inoculum was increased by one log, transmission chains were only broken in one out of three replicates. If the same experiment was performed with a short cohousing period (24 hours), transmission chains were blocked efficiently with low and high inocula; at the low inoculum dose, even transmission to the first recipient was blocked in two out of three replicates. These different scenarios may be similar to situations that humans experience during the influenza season as well. The short exposure experiment may resemble short contacts with infected individuals, e.g., in public transport, during a dinner or at work. The long exposure is perhaps akin to exposure to infected family members within a household. The low and high inocula perhaps resemble close contact without a mask versus less close contact or masking. Irrespectively, in three out of four scenarios, mucosal immunity to NA was able to break transmission chains and similar immunity in the human population may restrict influenza virus circulation during the influenza season to a large degree. We cannot exclude that recombinant HA would have the same effect. Indeed, it is likely that a recombinant HA vaccine administered the same way would perform well. However, antigenic drift may affect HA more than NA and we therefore think, based on the data presented here and a large number of studies by us and others that show benefits of NA-based immunity, further (clinical) development of NA-based mucosal vaccines is warranted (17, 20).

## Acknowledgements

This work was supported in part by the National Institute of Allergy and Infectious Disease (NIAID) grant AI117287, the Collaborative Influenza Vaccine Innovation Centers (CIVIC) contract 75N93019C00051 (F.K.) and the NIAID Centers of Excellence for Influenza Research and Response (CEIRR) contract 75N93021C00014 (F.K.).

## Data availability statement

Data will be made publicly available upon publication and upon request for peer review.

